# A Tethered Ligand Assay to Probe the SARS-CoV-2 ACE2 Interaction under Constant Force

**DOI:** 10.1101/2020.09.27.315796

**Authors:** Magnus S. Bauer, Sophia Gruber, Lukas F. Milles, Thomas Nicolaus, Leonard C. Schendel, Hermann E. Gaub, Jan Lipfert

**Affiliations:** Department of Physics and Center for NanoScience (CeNS), LMU Munich, Amalienstrasse 54, 80799 Munich, Germany; Department of Biochemistry, University of Washington, Seattle, WA 98195, USA; Institute for Protein Design, University of Washington, Seattle, WA 98195, USA

## Abstract

The current COVID-19 pandemic has a devastating global impact and is caused by the SARS-CoV-2 virus. SARS-CoV-2 attaches to human host cells through interaction of its receptor binding domain (RBD) located on the viral Spike (S) glycoprotein with angiotensin converting enzyme-2 (ACE2) on the surface of host cells. RBD binding to ACE2 is a critical first step in SARS-CoV-2 infection. Viral attachment occurs in dynamic environments where forces act on the binding partners and multivalent interactions play central roles, creating an urgent need for assays that can quantitate SARS-CoV-2 interactions with ACE2 under mechanical load and in defined geometries. Here, we introduce a tethered ligand assay that comprises the RBD and the ACE2 ectodomain joined by a flexible peptide linker. Using specific molecular handles, we tether the fusion proteins between a functionalized flow cell surface and magnetic beads in magnetic tweezers. We observe repeated interactions of RBD and ACE2 under constant loads and can fully quantify the force dependence and kinetics of the binding interaction. Our results suggest that the SARS-CoV-2 ACE2 interaction has higher mechanical stability, a larger free energy of binding, and a lower off-rate than that of SARS-CoV-1, the causative agents of the 2002-2004 SARS outbreak. In the absence of force, the SARS-CoV-2 RBD rapidly (within ≤1 ms) engages the ACE2 receptor if held in close proximity and remains bound to ACE2 for 400-800 s, much longer than what has been reported for other viruses engaging their cellular receptors. We anticipate that our assay will be a powerful tool investigate the roles of mutations in the RBD that might alter the infectivity of the virus and to test the modes of action of neutralizing antibodies and other agents designed to block RBD binding to ACE2 that are currently developed as potential COVID-19 therapeutics.

## INTRODUCTION

Severe acute respiratory syndrome-corona virus-2 (SARS-CoV-2) is the causative agent of coronavirus disease-2019 (COVID-19), which emerged in late 2019. SARS-CoV-2 particles carry ∼100 copies of the trimeric viral glycoprotein Spike (S) on their surface^1^, giving the appearance of an eponymous corona around the virus. Like SARS-CoV-1, which caused an outbreak in 2002-2004, SARS-CoV-2 attaches to human host cells by S binding to angiotensin converting enzyme-2 (ACE2)^2, 3, 4, 5, 6^ (**Fig. 1A,B**). Specifically, each S trimer carries receptor binding domains (RBD) at the tip of the three S1 domain that can bind to ACE2 (**Fig. 1A,B**). Binding of the virus to host cells occurs in dynamic environments^7, 8^ where external forces act on the virus particle. In particular in the upper and lower respiratory tract, coughing, sneezing, and mucus clearance exert mechanical forces^9, 10^ that the virus must withstand for productive infection. In addition, standard binding assays suggest dissociation constants for isolated SARS-CoV-2 RBD binding to ACE2 in solution in the range *K*_d_ ∼ 1-100 nM (**Supplementary Table 1**), while the estimated concentration of S *in vivo* is ∼1 pM, based on 7·10^6^ viral copies per ml sputum^7^ and 100 S proteins per virus^1^ − orders of magnitude lower than the measured *K*_d_. To enhance both avidity and force stability, SARS-CoV-2 attachment to host cells very likely involves multivalent interactions. The homotrimeric nature of S, combined with the dense coverage of the viral capsid surface by S trimers^1^ and the observation that ACE2 clusters on the apical site of cells^3^ imply a high local density of binding partners. Consequently, an initial binding event could rapidly lead to further engagement of additional ligand-receptor pairs^11^ as has been suggested for a number of other viruses, including influenza, rabies, and HIV^12, 13, 14^. Stable binding of S to ACE2 enables further downstream events such as cleavage of S by furine or TMPRSS2 proteases^5, 11, 15^, triggering conformational changes, and ultimately fusion with the cell membrane and cellular entry.

**Figure 1.**
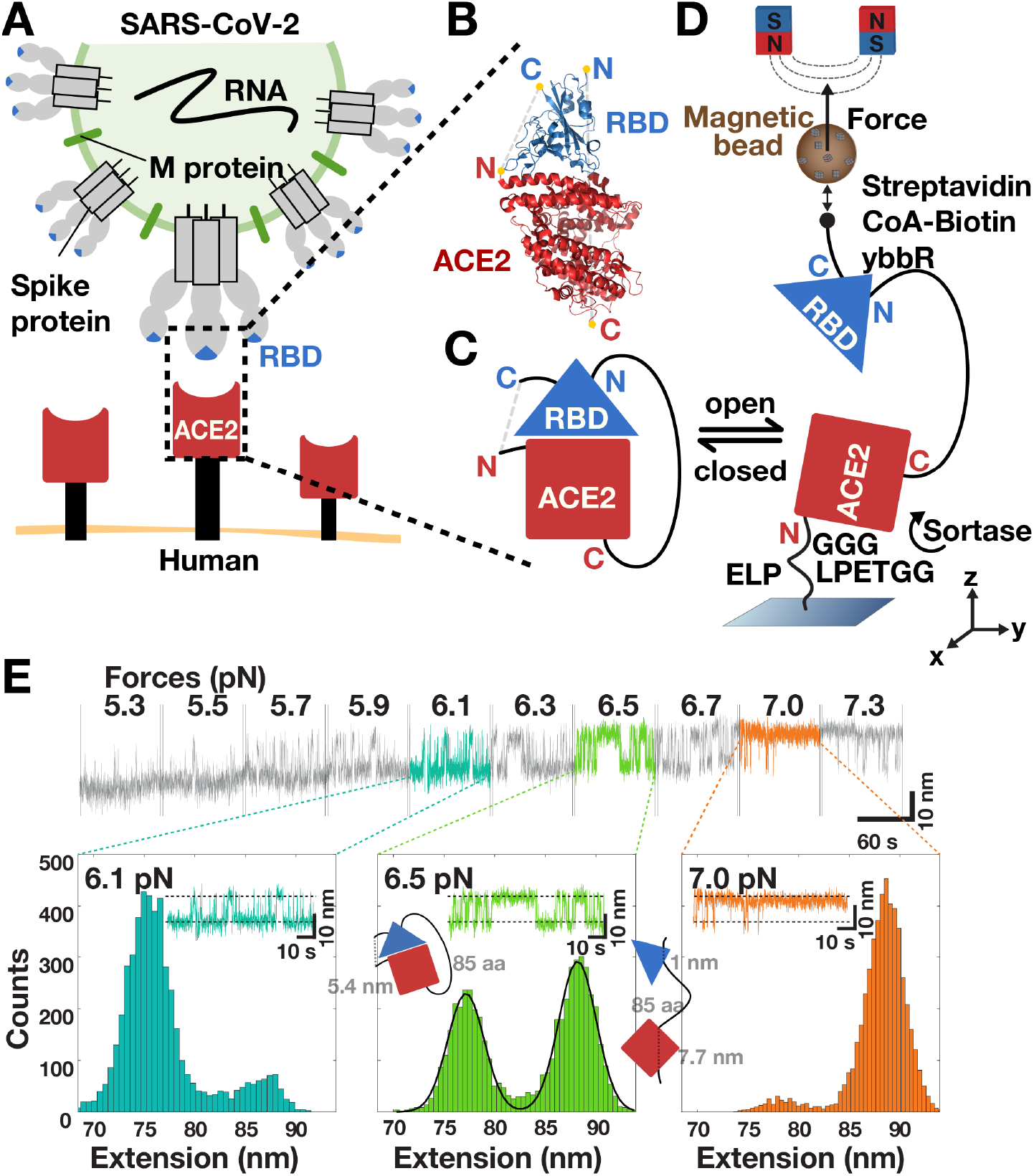
A tethered ligand assay probes the SARS-CoV-2 ACE2 interaction in magnetic tweezers. **A** Schematic rendering of SARS-CoV-2 (top) binding to human cells (bottom). The virus binds via its RBD (blue triangle) located at the tip of the S1 domain in each copy of the S protein trimer and engages the ectodomain of ACE2 (red rectangle) that is anchored to the cell membrane by its transmembrane domain (black rectangle). **B** Structure of the SARS-CoV-2 RBD bound to ACE2 (rendered from PDB entry 6M0J^32^) with the N- and C-termini of both proteins highlighted in yellow. **C** Scheme of the fusion protein receptor ligand construct. Shown is the variant N-terminus-ACE2-linker-RBD-C-terminus. **D** Schematic of the MT tethered receptor ligand assay. The fusion protein construct shown in C is attached via an ELP linker to a flow cell surface and via biotin-streptavidin to magnetic beads (for details of the molecular handles and protocol used for attachment see Materials and Methods). Permanent magnets mounted above the flow cell are used to apply calibrated stretching forces to the tether. **E** Extension time traces of a ACE2-linker-SARS-CoV-2 RBD fusion construct at different levels of applied force (indicated above the trace segments). Stochastic transition between a lower and a higher extension are observed that systematically shift to the higher extension state with increasing force. The overall shift in extension from plateau to plateau is due to the stretching response of the tether. Extension histograms (bottom) are shown for the three plateaus highlighted in color and reveal two distinct peaks. The two peaks are well described by a double Gaussian fit (solid line, middle panel) and correspond to the open and closed state of the tethered receptor ligand pair (shown schematically as an inset).

The SARS-CoV-2 S protein and its interaction with ACE2 have been the target of intense research activity, as they are critical in the first steps of SARS-CoV-2 infection and constitute a major drug target in the current search for treatments of COVID-19. Further, differences in binding between ACE2 and the SARS-CoV-1 and SARS-CoV-2 RBDs have been linked to the different observed patterns in lower and upper respiratory tract infections by the two viruses^5^. Despite its importance, many questions about RBD ACE2 interactions, in particular about their stability under constant external force, are unresolved. Consequently, there is an urgent need for assays that can probe the affinity and kinetics of the interaction under controlled external forces and that can mimic the effect of multivalent interactions *in vivo* by positioning the ligand-receptor pair in spatial proximity at an effective concentration much higher than in solution-based methods.

Here we present a tethered ligand assay to determine RBD interactions with ACE2 at the single-molecule level subject to defined levels of applied force. Our assay utilizes fusion protein constructs comprising of SARS-CoV-1 or SARS-CoV-2 RBD and human ACE2 joined by flexible peptide linkers (**Fig. 1B,C**). We hold our tethered receptor ligand constructs under precisely controlled and constant external force in magnetic tweezers (MT)^16, 17^ (**Fig. 1D**). Tethered ligand assays have provided insights into von Willebrand Factor binding to platelets^18, 19^, force-sensing of the cytoskeletal protein filamin^20^, rapamycin-mediated association between FKBP12 and FRB^21^, and protein-histone interactions^22^. Their key advantage is that they allow observation of repeated interactions of the same binding partners that are held in spatial proximity under mechanical control. Therefore, they can provide information about affinity, avidity, on- and off-rates, and mechanical stability. Measuring at the single-molecule level naturally provides access to kinetics and molecular heterogeneity. Using the tethered ligand assay, we compare the stability of the SARS-CoV-1 and SARS-CoV-2 RBD ACE2 interactions under mechanical load, measure the on- and off-rates, and extrapolate to the thermodynamic stability at zero load. Our assay gives direct access to binding rates of ligand-receptor pairs held in spatial proximity and we anticipate that it will be a powerful tool to assess the mode of action of potential therapeutic agents (such as small molecules^23^, neutralizing antibodies^24, 25^, nanobodies^26, 27, 28^, or designer proteins^29, 30^) that interfere with S binding to ACE2.

## RESULTS

### A tethered ligand assay to probe SARS-CoV RBD interactions with ACE2 in MT

We designed tethered ligand fusion proteins that consist of the SARS-CoV-1 or SARS-CoV-2 RBD and the ectodomain of human ACE2 joined by flexible peptide linkers (**Fig. 1B,C**). Protein constructs were designed based on the available crystal structures^31, 32^ of the SARS-CoV-1 or SARS-CoV-2 RBDs in complex with human ACE2 and carry short peptide tags at their termini for attachment in the MT (**Fig. 1D;** for details see **Materials and Methods**). Protein constructs were coupled covalently to the flow cell surface via elastin-like polypeptide (ELP) linkers^33^ and to magnetic beads via the biotin-streptavidin linkage. Tethering proteins via ELP linkers in the MT enables parallel measurements of multiple molecules over extended periods of time (hours to weeks) at precisely controlled forces^34^. In the MT, bead positions and, therefore, tether extensions are tracked by video microscopy in (x,y,z) with ∼1 nm spatial resolution and up to kHz frame rates^35, 36, 37^.

### Observation of RBD ACE2 interactions under force in MT

After tethering the fusion protein constructs in the MT, we subjected the protein tethers to different levels of constant force and recorded time traces of tether extensions (**Fig. 1E**). At forces in the range of 2-7 pN, we observed systematic transitions in the extension traces, with jumps between a high extension “open” and low extension “closed” state (**Fig. 1E**). The transitions systematically changed with applied force: At low forces, the system is predominantly in the closed state, while increasing force systematically increases the time spent in the open state. Histograms of the tether extension revealed two clearly separated peaks (**Fig. 1E**, bottom and **Fig. 2A,D**). By setting thresholds at the minima between the extension peaks, we defined populations in the open and closed states. The fraction in the open state systematically increases with increasing force (**Fig. 2B,E**; symbols) following a sigmoidal force dependence. The data are well described by a simple two-state model (**Fig. 2B,E**; solid line) where the free energy difference between the two states depends linearly on applied force *F*, i.e. Δ*G =* Δ*G*_0_ *–F·*Δ*z*, such that the fraction in the open state is given by

**Figure 2.**
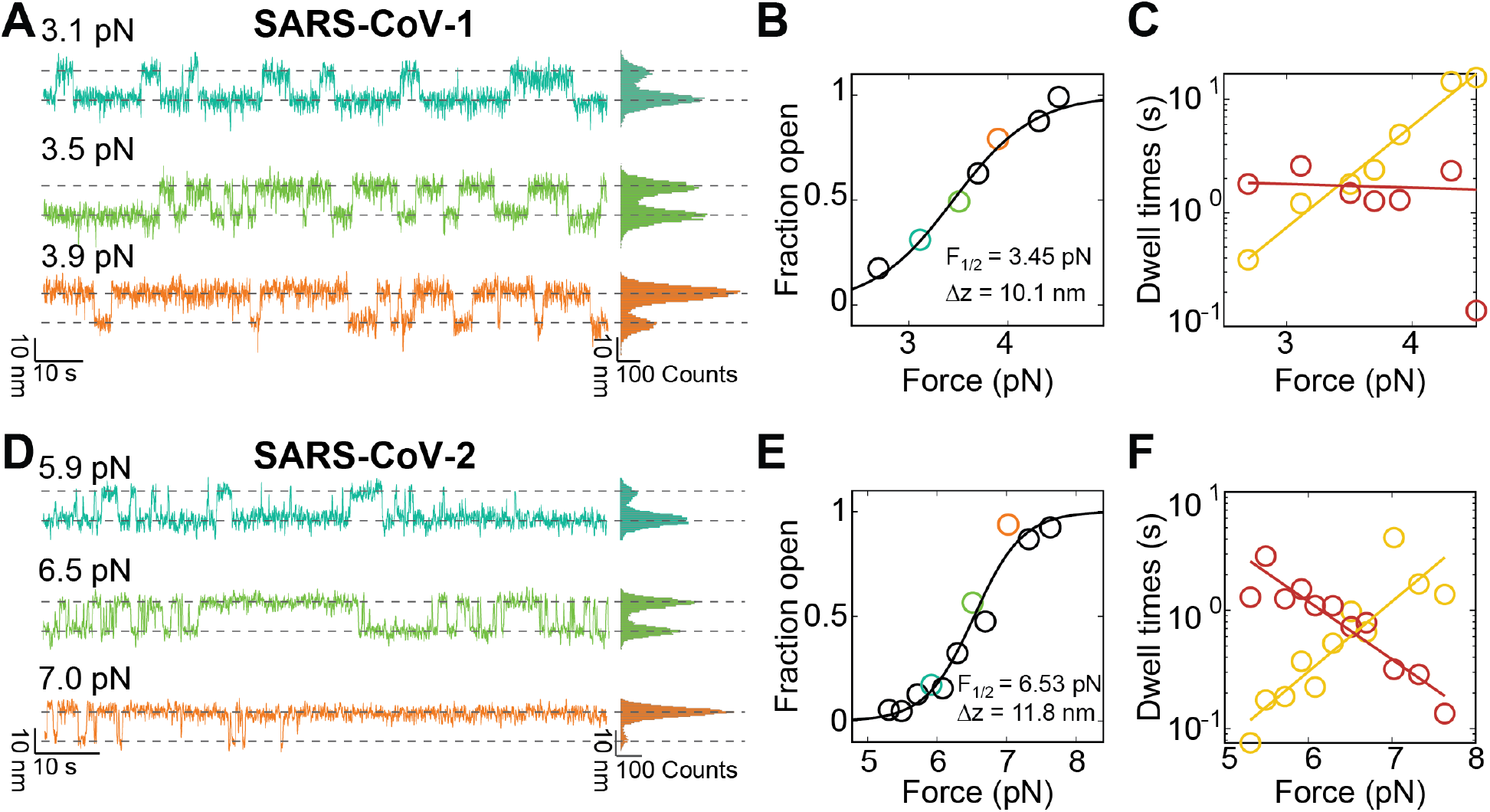
Comparison of mechanical stability and kinetics of ACE2 binding to SARS-CoV-1 and SARS-CoV-2 RBDs. **A** Extension time traces at different levels of applied force for the ACE2-linker-SARS-CoV-1 RBD fusion construct reveal systematic transitions between a low extension closed state and a high extension open state. Increasing force increases the fraction of time spent in the higher extension open conformations. **B** Quantification of the fraction open from extension time traces as a function of applied force (symbols; points determined from the traces in panel A are shown with matching color codes). The black line is a fit of the model shown in Equation 1. Fitting parameters are shown as an inset. **C** Mean dwell times in the open (yellow) and closed (dark red) states as a function of applied force. Mean dwell times were determined from maximum likelihood fits of a single exponential to the dwell time distributions. The solid lines are linear fits to the logarithm of the rate, i.e. to the model shown in Equation 2. **D,E,F** Same as panels A-C, but for the ACE2-linker-SARS-CoV-2 RBD construct.

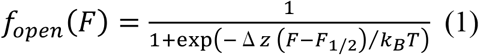

where *k*_*B*_ is Boltzmann’s constant, *T* the absolute temperature, and *F*_1/2_ and Δ*z* are fitting parameters that represent the midpoint force, where the system spends half of the time in the open and half of the time in the closed conformation, and the distance between the two states along the pulling direction, respectively. The free energy difference at zero force is given by Δ*G*_0_ = *F*_1/2_*·*Δ*z* and provides a direct measure of the stability of the binding interface.

From fits to the data for the construct ACE2-linker-SARS-CoV-2 RBD (**Fig. 2E**), we found *F*_1/2_ = 5.7 ± 1.2 pN and Δ*z =* 12.0 ± 2.2 nm, and, therefore, Δ*G*_0_ = *F*_1/2_*·*Δ*z* = 10.1 ± 2.8 kcal/mol (data are the mean and standard deviation from fits to biological repeats; see **Table 1** for a summary of all fitted parameters). The value of Δ*z* determined from fits of Equation 1 is in excellent agreement with the distance between the open and closed states Δ*z*_G_ = 13.0 ± 2.1 nm determined from fitting two Gaussians to the extension histograms at the equilibrium force *F*_1/2_ and evaluating the distance between the fitted center positions. The observed Δ*z* is also in agreement with the expected extension change of ≈ 13.4 nm, based on the crystal structure^32^ (PDB code 6M0J) assuming that the individual domains (ACE2 ectodomain and RBD) are rigid and remain folded in the open conformation and taking into account the stretching elasticity of the 85 amino acid (aa) protein linker using the the worm-like chain (WLC) model^34, 38, 39^ with a bending persistence length of *L*_p_ = 0.4 nm and contour length of *L*_c_ = 0.4 nm/aa (**Supplementary Fig. S1**).

**Table 1.** Interactions parameters for ACE2 and SARS-CoV-1 or SARS-CoV-2 RBD determined using the tethered ligand assay. Data are the mean and standard deviation from *N* = 6, 4, 9, and 7 molecules, respectively.

|  | **SARS-CoV-2:**<br>**ACE2-linker-**<br>**RBD**<br>**(85 aa linker)** | **SARS-CoV-2:**<br>**RBD-linker-**<br>**ACE2**<br>**(85 aa linker)** | **SARS-CoV-1:**<br>**ACE2-linker-**<br>**RBD**<br>**(85 aa linker)** | **SARS-CoV-1:**<br>**ACE2-linker-**<br>**RBD**<br>**(115 aa linker)** |
| --- | --- | --- | --- | --- |
| $F_{1/2}$ | $5.7 \pm 1.2$ pN | $4.2 \pm 1.0$ pN | $3.3 \pm 0.4$ pN | $3.5 \pm 0.7$ pN |
| $\Delta z$ (from fit of Equation 1) | $12.0 \pm 2.2$ nm | $11.2 \pm 0.8$ nm | $9.4 \pm 1.9$ nm | $14.0 \pm 2.9$ nm |
| $\Delta z_G$ (from fit of two Gaussians) | $13.0 \pm 2.1$ nm | $10.9 \pm 3.0$ nm | $11.8 \pm 1.2$ nm | $12.4 \pm 3.9$ nm |
| $\Delta G_0 (= \Delta z \cdot F_{1/2})$ | $10.1 \pm 2.8$ kcal/mol | $6.6 \pm 1.7$ kcal/mol | $4.4 \pm 1.0$ kcal/mol | $6.9 \pm 1.9$ kcal/mol |
| $\tau_{0,open}$ | $(2.4 \pm 2.8) \cdot 10^{-4}$ s | $(6.4 \pm 7.5) \cdot 10^{-4}$ s | $(8.4 \pm 7.0) \cdot 10^{-3}$ s | $(2.7 \pm 1.9) \cdot 10^{-4}$ s |
| $\tau_{0,closed}$ | $435 \pm 493$ s | $797 \pm 907$ s | $22 \pm 49$ s | $14 \pm 7$ s |
| $\Delta G_{0,\tau} (= k_B T \cdot \log(\tau_{0,open}/\tau_{0,closed}))$ | $8.8 \pm 2.1$ kcal/mol | $8.4 \pm 2.0$ kcal/mol | $4.1 \pm 1.2$ kcal/mol | $6.5 \pm 0.4$ kcal/mol |
| $\Delta z_{open}$ | $6.4 \pm 1.3$ nm | $9.4 \pm 5.4$ nm | $6.8 \pm 1.2$ nm | $9.4 \pm 1.3$ nm |
| $\Delta z_{closed}$ | $4.2 \pm 1.4$ nm | $4.7 \pm 3.2$ nm | $1.9 \pm 2.3$ nm | $3.9 \pm 1.5$ nm |

A construct using the same 85 aa linker and same attachment geometry, but the SARS-CoV-1 RBD instead of SARS-CoV-2 RBD, showed a qualitatively very similar behavior (**Fig. 2A,B**), with stochastic transitions between an open and a closed conformation. From fits of Equations 1, we found *F*_1/2_ = 3.3 ± 0.4 pN and Δ*z =* 9.4 ± 1.9 nm and thus Δ*G*_0_ = 4.4 ± 1.0 kcal/mol for SARS-CoV-1 (**Table 1**). The midpoint force and binding energy are, therefore, approximately two-fold lower for SARS-CoV-1 RBD interacting with ACE2 compared to SARS-CoV-2 using the same linker and a very similar overall geometry. The length increment Δ*z* determined from fits of Equation 1 is again, within experimental error, in agreement with the value determined from fitting two Gaussians to the extension histogram near the midpoint of the transition (Δ*z*_G_ = 11.8 ± 1.2 nm at *F*_1/2_) and with the expected extension change of ≈ 12.1 nm taking into account the crystal structure of the SARS-CoV-1 RBD bound to ACE2 ^31^ (PDB code 2AJF). The slightly shorter extension increment upon opening for the SARS-CoV-1 construct compared to SARS-CoV-2, despite using the same 85 aa linker and a very similar crystallographic geometry is mostly due to the smaller extension of the WLC at the lower midpoint force for SARS-CoV-1. Control measurements for the same ACE2-SARS-CoV-1 RBD construct with a 115 aa instead of 85 aa linker show a larger length increment Δ*z* = 14.0 ± 2.9 nm upon opening, consistent with the expectation of ≈ 15.1 nm from a longer linker and again with good agreement between the Δ*z* value fitted from Equation 1 and Δ*z*_G_ from Gaussian fits of the extension histogram (**Table 1**).

As an additional control measurement to test for possible influences of the linker insertion and coupling geometry, we used an inverted geometry with force applied to the N-terminus of the SARS-CoV-2 RBD and to the C-terminus of ACE2, again with an 85 aa linker. The inverted construct showed similar stochastic transitions between an open and a closed state (**Supplementary Fig. S2**). We found *F*_1/2_ = 4.2 ± 1.0 pN and Δ*z =* 11.2 ± 0.8 nm from fits of Equation 1, again in excellent in agreement with Δ*z*_G_ *=* 10.9 ± 3.0 nm. The predicted length change from the crystal structure is ≈ 6.2 nm, still in rough agreement but slightly shorter than the experimentally determined value, while the prediction for the opposite geometry was close to or slightly longer than what was determined from the extension traces. The overall good agreement between predicted and measured length increments upon opening of the complexes and the fact that the deviations have the opposite sign for the two different tethered ligand geometries strongly suggest that the RBD and ACE2 ectodomain remain folded in the open conformations. Significant unfolding of the domains upon opening of the complex would increase the observed length increment compared to the predictions that assume folded domains and lead to systematically larger measured compared to predicted Δ*z* values. We note that some residues are not resolved in the crystal structure and, therefore, not taken into account in our prediction (**Supplementary Fig. S1**). The observed deviations between predicted and measured Δ*z* values would be consistent with the unresolved residues at the RBD C-terminus becoming part of the flexible linker and the missing residues at the N-terminus remaining folded as part of the RBD. Taken together, the MT data show that our tethered ligand assay can systematically probe RBD ACE2 binding as a function of applied force and enables faithful quantitation of the mechanostability and thermodynamics of the interactions.

### The tethered ligand assay gives access to ACE2 RBD binding kinetics under force

In addition to providing information on the binding equilibrium, the tethered ligand assay probes the binding kinetics under force. Analyzing the extension-time traces using the same threshold that was used to determine the fraction open vs. *F*, we identify dwell times in the open and closed states (**Supplementary Fig. S3A,B**). We find that the dwell times in the open and closed states are exponentially distributed (**Supplementary Fig. S3C,D**). The mean dwell times in the closed state decrease with increasing force, while the mean dwell times in the open state increase with increasing force (**Fig. 2C,F**). The dependencies of the mean dwell times on applied force *F* are well described by exponential, Arrhenius-like relationships ^40^

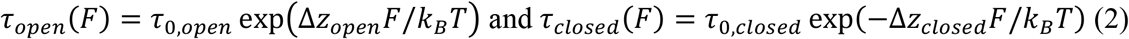

where the fitting parameters τ_0,open_ and τ_0,closed_ are the lifetimes of the open and closed conformation in the absence of force and Δ*z*_open_ and Δ*z*_closed_ are the distances to the transition state along the pulling direction.

For all constructs measured, the sum Δ*z*_open_ + Δ*z*_closed_ is equal, within experimental error, to the total distance between the open and closed conformations Δ*z* (**Table 1**), providing a consistency check between the equilibrium and kinetic analyses. The parameters Δ*z*_open_ and Δ*z*_closed_ quantify the force-dependencies of the lifetimes of the respective states and the slopes in the log(τ_open/closed_) vs. *F* plots (**Fig. 2 C,F**) are given by Δ*z*_open/closed_ / *k*_*B*_*T*. For all tethered ligand constructs investigated, Δ*z*_closed_ is smaller than Δ*z*_open_ (by approximately a factor of ∼2), i.e. opening of the bound complex is less force sensitive than rebinding from the open conformation. The different force sensitivities can be rationalized from the underlying molecular processes: The closed complexes feature protein-protein interfaces that will break over relatively short distances; in contrast, the open conformations involve flexible peptide linkers that make rebinding from the open states more force dependent.

The extrapolated lifetimes at zero force of the closed conformations τ_0,closed_ are in the range of 400-800 s for the SARS-CoV-2 and ∼20 s for SARS-CoV-1. In comparison, the lifetimes of the open states in the absence of load τ_0,open_ are much shorter, in the range of ∼1 ms (**Table 1**). The extrapolated lifetimes at zero force provide an alternative route to computing the free energy difference between the open and closed conformations at *F* = 0, which is given by ΔG_0,tau_ = *k*_*B*_*T* · log(τ_0,open_/τ_0,closed_). For all constructs, we find excellent agreement, within experimental error, between the free energy differences ΔG_0,tau_ determined from the extrapolated lifetimes and the values Δ*G*_0_ = *F*_1/2_*·*Δ*z* from Equation 1 (**Table 1**). The close agreement of the ΔG_0,tau_ and Δ*G*_0_ values provides another consistency check between the kinetic and equilibrium analyses. The results show that our tethered ligand assay can yield consistent and complementary information both on the binding equilibrium and on the interaction kinetics under external force.

### Quantitative comparison of tethered ligand data to free solution binding assays

Traditional binding assays measure the interaction of binding partners in free solution. In contrast, the tethered ligand assay probes binding between receptor-ligand pairs held in proximity and under external force. While the situation *in vivo* is even more complex, the tethered ligand assay mimics the multivalent interactions that likely occur between viral particles with multiple trimeric S complexes and the apical surface of cells where multiple binding partners are in spatial proximity. To compare tethered ligand measurements to traditional binding assays, it is important to consider the differences between tethered ligand-receptor systems and cases with binding partners in free solution. The free energies ΔG_0_ (or ΔG_0,tau_) determined in our assay measure the stability of the bound complex with respect to the open state with the ligand tethered. Consequently, ΔG_0_ will in general depend on the length of the linker and the tethering geometry, as we clearly observe experimentally: For the same set of binding partners, we find significantly different values for ΔG_0_ for different tethering geometries. For example, we can compare ACE2 binding to the SARS-CoV-2 RBD in the two different tethering geometries (10.1 ± 2.8 kcal/mol vs. 6.6 ± 1.7 kcal/mol; *p* = 0.04 from a two-sample t-test) or the SARS-CoV-1 data for the 85 or 115 aa tethers (4.4 ± 1.0 kcal/mol vs. 6.8 ± 1.0 kcal/mol; *p =* 0.004). In contrast, binding assays with the binding partners in free solution are sensitive to the free energy difference between the bound complex and the ligand and receptor in solution, which depends on the solution concentrations.

To compare the two scenarios, it is useful to consider the problem in terms of lifetimes or, equivalently, (on- and off-) rates^19^. The lifetime of the bound complex in the tethered ligand system τ_0,closed_ (= 1/*k*_0,off_) has units of seconds and can be directly compared to the binding lifetimes (or solution off-rates *k*_sol,off_) measured in bulk binding assays. The lifetime of the open conformation in the tethered ligand assay τ_0,open_ (= 1/k_0,on_) also has units of seconds, but can not be directly compared to solution on-rates, since for a bimolecular reaction the solution on-rate *k*_sol,on_ has unit of M^−1^ s^−1^ and depends on concentration. To relate the two quantities, one can introduce an effective concentration ^19, 41, 42^ of the tethered ligand *c*_eff_ such that *k*_sol,on_ = *k*_0,on_ / *c*_eff_.

We can quantitatively relate our results to studies that have reported equilibrium dissociation constants and rates for the ACE2 interactions with SARS-CoV-1 and SARS-CoV-2 using traditional binding assays (for an overview see **Supplementary Table 1**). While the values reported in the literature vary significantly, likely due to the different experimental methods and sample preparation strategies used, clear and consistent trends can be identified. The lifetimes of the closed complex determined in our assay correspond to rates of *k*_0,off_ ∼ 5·10^−2^ s^−1^ for SARS-CoV-1 and ∼2·10^−3^ s^−1^ for SARS-CoV-2, well within the ranges of reported *k*_sol,off_ values in the literature^25, 32, 43, 44, 45^ (**Supplementary Table 1**). Our value for the off-rate of SARS-CoV-2 RBD bound to ACE2 is also in reasonable agreement with the value of (8 ± 5)·10^−3^ s^−1^ extrapolated from AFM force spectroscopy experiments^46^. A clear trend is that the off-rate for SARS-CoV-2 is smaller than for SARS-CoV-1, by about one order-of-magnitude, indicating a longer lived bound complex for the new SARS variant. In contrast, for the on-rates most solution binding assays report similar values for the two SARS variants, in the range of *k*_sol,off_ ∼10^5^ M^−1^ s^−1^. Our tethered ligand assay also found similar unimolecular on-rates for the two SARS variants, similar to ∼10^3^ s^−1^, implying an effective concentration of *c*_eff_ = *k*_0,on_ / *k*_sol,on_ ∼ 10 mM. This effective concentration is in the range of concentrations found for other tethered ligand protein systems^19, 42, 47^ and can be understood as the apparent concentration of one molecule in a sphere of ∼4 nm radius, a distance close to distances to the transition states determined from the data under force and to the approximate mean square end-to-end distance in solution for a 85 aa peptide.

Taken together, we find that in the absence of applied force, the SARS-CoV-2 RBD remains bound to ACE2 for ∼400-800 s, consistent with traditional binding assays and at least 10-fold longer than the lifetime of the SARS-CoV-1 RBD interaction with ACE2. The time scale for binding in free solution is concentration dependent, but for the situation that the binding partners are held in close proximity, we observe rapid (re-)binding within <1 ms in the absence of force for both SARS variants.

## CONCLUSION

We have developed a tethered ligand assay to probe SARS-CoV RBD interactions with ACE2 under precisely controlled levels of applied force. Our approach provides quantitative information about both binding equilibrium and kinetics. We find that a single SARS-CoV-2 RBD ACE2 interaction can withstand constant loads up to 5 pN (at least for ∼minutes time scales). We observe that the SARS-CoV-2 RBD interaction has a ∼2-fold higher force stability than SARS-CoV-1 using a similar tethering geometry. The higher force stability of SARS-CoV-2 compared to SARS-CoV-1 observed in our assay at constant force is qualitatively in line with recent data from AFM force spectroscopy at constant loading rate^48^. The higher force stability of SARS-CoV-2 engaging ACE2 might contribute to fact that SARS-CoV-2 more frequently infects the upper respiratory tract in addition to deep lung tissue compared to the 2002 SARS variant^7, 49^.

We find that in the absence of applied force, the SARS-CoV-2 RBD remains bound to ACE2 for ∼400-800 s, which would provide a long time window for conformational rearrangements to engage additional RBD copies on the same S trimer^11^, to bind to additional S trimers, and to initiate proteolytic cleavage and downstream processes. Our measured lifetime of the initial RBD ACE2 interaction is much longer than the values < 1 s reported for influenza, rabies, or HIV viruses engaging their cellular receptors measured by AFM or optical tweezers force spectroscopy^12, 13, 50, 51^, which might contribute to SARS-CoV-2 higher infectivity. For influenza, rabies, and HIV multivalent interactions of the virus with its host cell have been suggested to play important roles^12, 13, 14^. Our data suggest that if held in close proximity, SARS-CoV RBDs can engage ACE2 rapidly, within τ_0,open_ ∼ 1 ms. While our assay is different from the situation *in vivo*, the tethered ligand mimics the effect of pre-formed interactions by a subset of the RBDs in the S trimer or by neighboring S trimers, which suggests that multivalent interactions between the virus and its host cell could form rapidly after an initial binding event, providing additional stability of the interaction. We estimate the concentration of S *in vivo* as ∼1 pM, based on 7·10^6^ viral copies in ml sputum^7^ and 100 S proteins per virus^1^. This estimated bulk protein concentration *in vivo* is much lower than the dissociation constants reported, which are in the range *K*_d_ ∼ 1-100 nM for the SARS-CoV-2 RBD binding to ACE2 and 10-fold lower affinity for SARS-CoV-1 (**Supplementary Table 1**), suggesting that multivalency might be critical for efficient viral binding. The rapid binding of RBDs held in proximity to ACE2 revealed by our assay might, therefore, be an important component of SARS-CoV-2 infections.

We anticipate that our tethered ligand assay will provide a powerful approach to investigate how the RBD ACE2 binding is blocked or altered by antibodies, nanobodies, or other drugs. In particular, the tethered ligand assay could go beyond standard bulk assays and reveal heterogeneity, include avidity effects, and determine drug residence times, in addition to affinities^21^. In addition, our approach could provide a tool to characterize emerging mutations of the viral S protein that alter binding or interfere with antibody recognition^24, 52^. Beyond the current COVID-19 pandemic, our assay provides a new method to probe cell-virus interactions^53^ and should be broadly applicable to other host-pathogen interactions.

## ACKNOWLEDGEMENTS

We thank Rafael C. Bernardi, David Dulin, Daniel Lietha, and Klaus Überla for helpful discussions. This study was supported by German Research Foundation Project 386143268, an EMBO long term fellowship to L.F.M. (ALTF 1047-2019), and the Physics Department of the LMU Munich.

## MATERIALS AND METHODS

### Cloning and Protein Construct Design

Constructs for ACE2-linker-RBD of SARS-CoV-1 were designed in SnapGene Version 4.2.11 (GSL Biotech LLC) based on a combination of the ACE2 sequence from Komatsu *et al*.^54^ available from GenBank under accession number AB046569 and the SARS-CoV-1 sequence from Marra *et al*.^55^ available from GenBank under accession number AY274119. The crystal structure by Li *et al*.^31^ available from the Protein Data Bank (PDB accession number 2AJF) was used as a structural reference. The linker sequence and tag placement was adapted from Milles *et al*.^56^. The linker sequence is a combination of two sequences available at the iGEM parts databank (accession numbers BBa_K404300, BBa_K243029). We used an analogous approach to design the fusion protein with the sequence of the RBD of SARS-CoV-2 from the sequence published by Wu e*t al*.^57^ available from GenBank under accession number MN908947. Reverse control constructs with C-terminal ACE2 were designed by reversing the order of the protein domains. A 6x histidine tag was added for purification. In addition, tags for specific pulling in magnetic tweezers were introduced: a triple glycine for sortase-catalyzed attachment on the N-terminus and a ybbR-tag on the C-terminus. In summary, the basic construct is built up as follows: MGGG-ACE2-linker-RBD-6xHIS-ybbR. All DNA and protein sequences are provided in the Supplementary Information.

The constructs were cloned using Gibson assembly from linear DNA fragments (GeneArt, ThermoFisher Scientific, Regensburg, Germany) containing the sequence of choice codon-optimized for expression in *E. coli* into a Thermo Scientific pT7CFE1-NHis-GST-CHA Vector (Product No. 88871). Control constructs with different sized linkers were obtained by blunt end cloning with either deleting or adding sequences to linker. Replication of DNA plasmids was obtained by transforming in DH5-Alpha Cells and running overnight cultures with 7 ml lysogeny broth with 50 µg/ml carbenicillin. Plasmids were harvested using a QIAprep® Spin Miniprep Kit (QIAGEN).

### In Vitro Protein Expression

Expression was conducted according to the manual of 1-Step Human High-Yield Mini *in vitro* translation (IVT) kit (Product No. 88891X) distributed by ThermoFisher Scientific (Pierce Biotechnology, Rockford, IL, USA). All components, except 5X dialysis buffer, were thawed on ice until completely thawed. 5X dialysis buffer was thawed for 15 minutes and 280 µl were diluted into 1120 µl nuclease-free water to obtain a 1X dialysis buffer. The dialysis device provided was placed into the dialysis buffer and kept at room temperature until it was filled with the expression mix.

For preparing the IVTT expression mix, 50 µl of the HeLa lysate was mixed with 10 µl of accessory proteins. After each pipetting step the solution was gently mixed by stiring with the pipette. Then the HeLa lysate and accessory proteins mix was incubated for 10 minutes. Afterwards, 20 µl of the reaction mix was added. Then 8 µl of the specifically cloned DNA (0.5 µg/µl) was added. The reaction mix was then topped off with 12 µl of nuclease-free water to obtain a total of 100 µl. This mix was briefly centrifuged at 10,000 g for 2 minutes. A small white pellet appeared. The supernatant was filled into the dialysis device placed in the 1X dialysis buffer. The entire reaction was then incubated for 16 h at 30°C under constant shaking at 700 rpm. For incubation and shaking an Eppendorf ThermoMixer with a 2 ml insert was used. After 16 h the expression mix was removed and stored in a protein low binding reaction tube on ice until further use.

### Protein Purification

Purification was conducted using HIS Mag Sepharose® Excel beads together with a MagRack™ 6 closely following the provided protocol. Bead slurry was mixed thoroughly by vortexing. 200 µl of homogenous beads were dispersed in a 1.5 ml protein low binding reaction tube. Afterwards the reaction tube was placed in the magnetic rack and the stock buffer was removed. Next, the beads were washed with 500 µl of HIS wash buffer (25 mM TRIS-HCl, 300 mM NaCl, 20 mM imidazol, 10% vol. glycerol, 0.25 % vol. Tween 20, pH 7.8). Expressed protein from IVTT was filled to 1000 µl with TRIS buffered saline (25 mM TRIS, 72 mM NaCl, 1 mM CaCl_2_, pH 7.2) and mixed with freshly washed beads. The mix was incubated in a shaker for 1 h at room temperature. Subsequently, the reaction tube was placed in the magnetic rack and the liquid was removed. The beads were washed three times with wash buffer keeping the total incubation time to less than 1 min. Remaining wash buffer was removed and 100 µl elution buffer (25 mM TRIS-HCl, 300 mM NaCl, 300 mM imidazol, 10% vol. glycerol, 0.25 % vol. Tween 20, pH 7.8) was added to wash protein off the beads. The bead elution buffer mix was then incubated for one minute with occasional gentle vortexing. Afterwards, the reaction tube was placed in the magnetic rack again to remove the eluted protein. This step was repeated for a second and third elution step. Buffer of the eluted protein was exchanged to TRIS buffered saline in 40k Zeba spin columns. Concentrations were determined photospectrometrically with a NanoDrop and aliquots were frozen in liquid nitrogen.

### Magnetic Tweezers Instrument

Measurements were were performed on a custom MT setup described previously^34, 37^. In the setup, molecules are tethered in a flow cell (FC; see next section); mounted above the FC is a pair of permanent magnets (5×5×5 mm^**3**^ each; W-05-N50-G, Supermagnete, Switzerland) in vertical configuration^17^. The distance between magnets and FC is controlled by a DC-motor (M-126.PD2; PI Physik Instrumente, Germany) and the FC is illuminated by an LED (69647, Lumitronix LED Technik GmbH, Germany). Using a 40x oil immersion objective (UPLFLN 40x, Olympus, Japan) and a CMOS sensor camera with 4096×3072 pixels (12M Falcon2, Teledyne Dalsa, Canada) a field of view of approximately 440 × 330 µm^**2**^ is imaged at a frame rate of 58 Hz. Images are transferred to a frame grabber (PCIe 1433; National Instruments, Austin, TX) and analyzed with an open-source tracking software^58, 59^. The tracking accuracy of our setup was determined to be ≈0.6 nm in (x, y) and ≈1.5 nm in z direction, as determined by tracking non-magnetic polystyrene beads, after baking them onto the flow cell surface. For creating the look-up table required for tracking the bead positions in z, the objective is mounted on a piezo stage (Pifoc P-726.1CD, PI Physik Instrumente). Force calibration was performed as described^60^ by analysis of the fluctuations of long DNA tethers. Importantly, for the small extension changes on the length scales of our protein tethers, the force stays constant to very good approximation (to better than 10^**−4**^ relative change). The largest source of force uncertainty is due to bead-to-bead variation, which is on the order of ≤ 10% for the beads used in this study^17, 61^.

### Flowcell Preparation and Magnetic Tweezers Measurements

Flowcells (FCs) were prepared as described previously^34^. Elastin-like polypeptide (ELP) linkers^33^ with a sortase motif at their C terminus and a single cysteine at their N terminus were coupled to aminosilanised glass slides via a small-molecule crosslinker with a thiol-reactive maleimide group^62^ (sulfosuccinimidyl 4-(N-maleimidomethyl)cyclohexane-1-carboxylate; sulfo-SMCC, ThermoFisher Scientific). 1 µm diameter polystyrene beads were baked onto the glass surface to serve as reference beads during the measurement. FCs were assembled from an ELP-functionalized bottom slide and an unfunctionalized glass slide with two holes (inlet and outlet) on either side serving as top slide. Both slides were separated by a layer of parafilm (Pechiney Plastic Packaging Inc., Chicago, IL, USA), which was cut out to form a 50 µl channel. FCs were incubated with 1% (v/v) casein solution (Sigma-Aldrich) for 3 to 4 h and flushed with 1 ml buffer (20 mM HEPES, 150 mM NaCl, 1 mM MgCl_2_, 1 mM CaCl_2_, pH 7.4).

CoA-biotin (New England Biolabs) was coupled to the ybbR-tag at the C-terminus of the fusion protein constructs in a 90 min bulk reaction in the presence of 4 µM sfp phosphopantetheinyl transferase^63^ and 100 mM MgCl_2_ at room temperature (≈ 22°C). Proteins were diluted to a final concentration of about 50 nM in 20 mM HEPES, 150 mM NaCl, 1 mM MgCl_2_, 1 mM CaCl_2_, pH 7.4. To couple the N-terminus of the fusion proteins carrying three glycines with the C-terminal LPETGG motif of the ELP-linkers, 100 µl of the protein mix was flushed into the FC and incubated for 25 min in the presence of 200 nM evolved pentamutant sortase A from *Staphylococcus aureus*^64, 65^. Unbound proteins were flushed out with 1 ml measurement buffer (20 mM HEPES, 150 mM NaCl, 1 mM MgCl_2_, 1 mM CaCl_2_, 0.1% (v/v) Tween-20, pH 7.4). Finally, commercially available streptavidin-coated paramagnetic beads (Dynabeads™ M-270 Streptavidin, Invitrogen) were added into the FC and incubated for 30 s before flushing out unbound beads with 1 ml measurement buffer. Receptor-ligand binding and unbinding under force was systematically investigated by subjecting the protein tethers to (90-120) s long plateaus of constant force, which was gradually increased in steps of 0.2 to 0.3 pN. All measurements were conducted at room temperature.

### Data Analysis

MT traces for analysis were selected on the basis of extension changes between an open and a closed state at forces between 1.5 and 7 pN, with a gradual shift towards an open state with increasing force. For each trace, (x,y)-fluctuations were checked to avoid inclusion of tethers that exhibit inter-bead or bead-surface interactions, which would also cause changes in x or y. Non-magnetic references beads were tracked simultaneously with magnetic beads and reference traces were subtracted for all measurements to correct for drift. Extension time traces were subjected to a 5-frame moving average smoothing to reduce noise. All analyses were performed with custom scripts in MATLAB.

## Supplementary Information

**Supplementary Figure S1.**
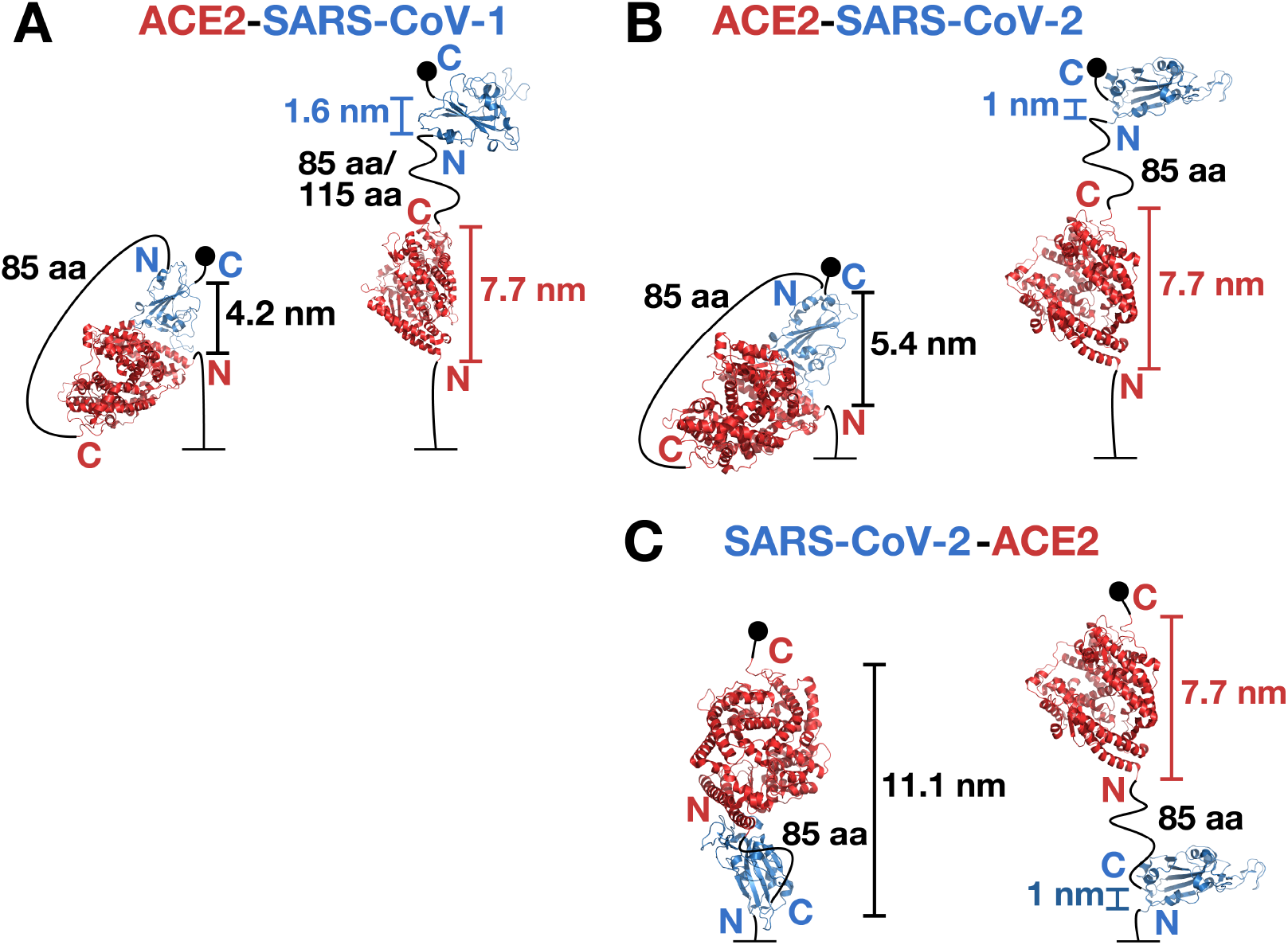
Estimation of the length increments Δz from crystal structures. For all constructs, schemes of the closed conformations are shown on the left and of the open conformations on the right. Closed conformations are based on the crystal structures^1, 2^ of RBD bound to ACE2, PDB accession codes 2AJF and 6M0J for SARS-CoV-1 and 2, respectively. Our simple estimates assume no deformations or flexibility of the crystal structures. For the closed conformations the distances between the N- and C-termini of the fusion constructs Δz_closed_ are determined from the crystal structure and indicated in the figure panels. The corresponding distances between the N- and C-termini of the fusion constructs in the open conformations Δz_open_ are estimated as follows: We assume that the RBD and ACE2 domains remain fully folded, but are free to rotate as indicated in the figure panels. The distance Δz_open_ is then given by the sum of the distances between the N- and C-termini of the individual domains (indicate in the figure panels) and the length of the ELP linker, which was estimated from the WLC model evaluated at the midpoint force *F*_1/2_ for each construct. The predicted extension increment Δz upon opening is given by Δz = Δz_open_ – Δz_closed_. **A** Estimate of the extension increment for the ACE2-linker-SARS-CoV-1 RBD construct. The extension of the 85 aa (115 aa) linker at *F*_1/2_ = 3.3 pN (3.5 pN) was computed to be 7.0 nm (10.0 nm). The predicted extension changes are, therefore, 12.1 nm and 15.1 nm, respectively. **B** Estimate of the extension increment for the ACE2-linker-SARS-CoV-2 RBD construct. The extension of the 85 aa linker at *F*_1/2_ = 5.7 pN was computed to be 10.1 nm. The predicted extension change is 13.4 nm. **C** Estimate of the extension increment for the SARS-CoV-2 RBD -linker-ACE2 construct. The extension of the 85 aa linker at *F*_1/2_ = 4.2 pN was computed to be 8.6 nm. The predicted extension change is 6.2 nm. We note that these simple estimates neglect the effect of several residues at the N- and C-termini of the RBD that are not resolved in the crystal structures (17 N-terminally and 10 C-terminally for SARS-CoV-2 and 17 N-terminally and 25 C-terminally for SARS-CoV-1).

**Supplementary Figure S2.**
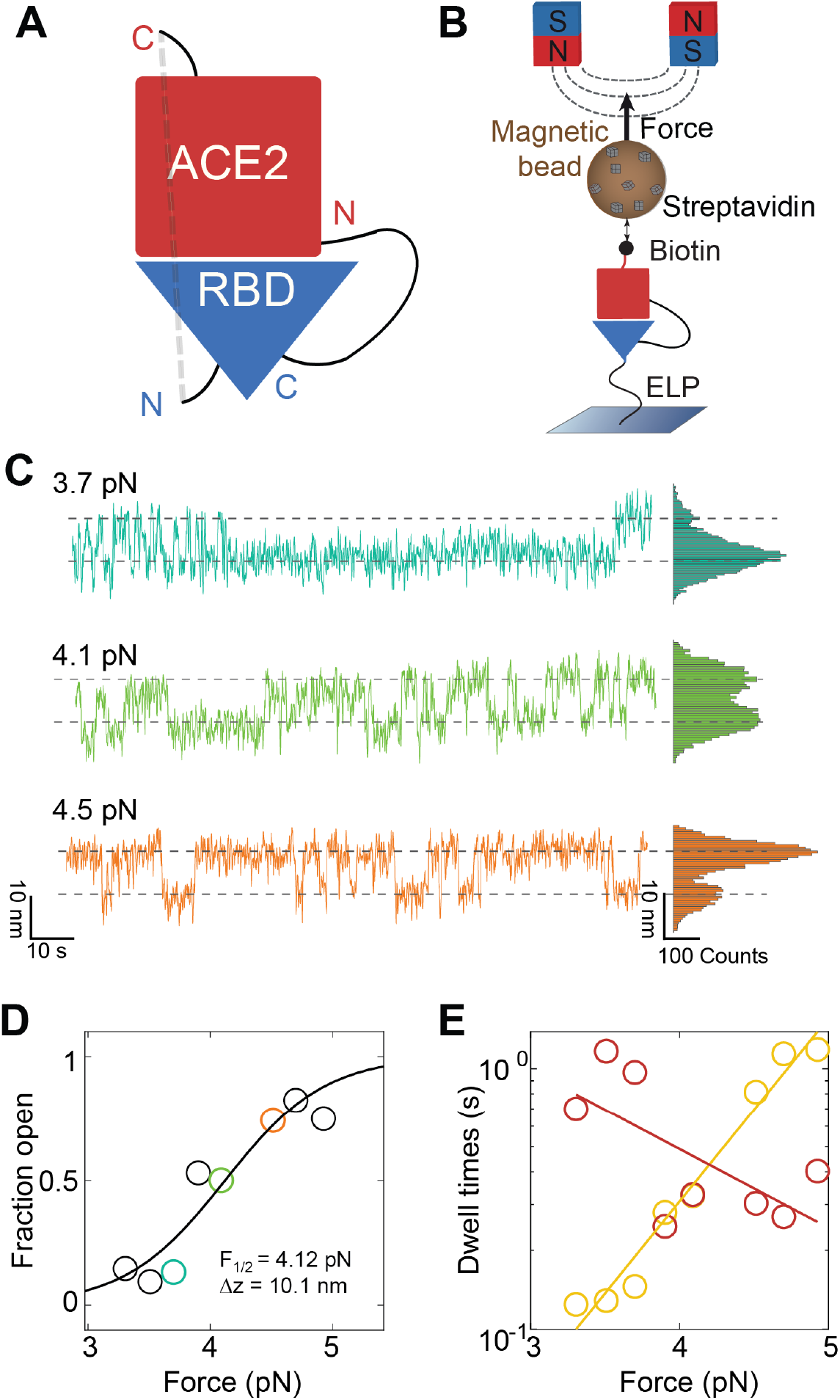
Mechanical stability and kinetics of the SARS-CoV-2 RBD ACE2 interaction using an inverted tethering geometry. **A** Schematic of the fusion protein construct used for measurements using an inverted geometry compared to the data shown in Fig. 1 and 2. Here, a flexible peptide linker connects the C-terminus of the SARS-CoV-2 RBD to the N-terminus ACE2 ectodomain. **B** Schematic of the alternative tethering geometry in the magnetic tweezers. The assay is identical to the scheme shown on Fig. 1C, except that now the ACE2 domain is attached via a ELP-linker to the flow cell surface and the RBD domain is coupled to biotin via the ybbR-tag and attached to streptavidin coated magnetic beads. **C** Extension time traces of the tether ligand construct with inverted geometry under different levels of constant force. The traces again reveal systematic transitions between low extension and high extension states, corresponding to the unbinding and (re-)binding of the RBD ACE2 interaction. **D** Fraction of time in the high extension open state as a function of applied force. The solid blue line is a fit of Equation 1 with fitting parameters indicated in the inset. **E** Mean dwell times in the open (yellow) and closed (dark red) states. Solid lines are fits of Equation 2.

**Supplementary Figure S3.**
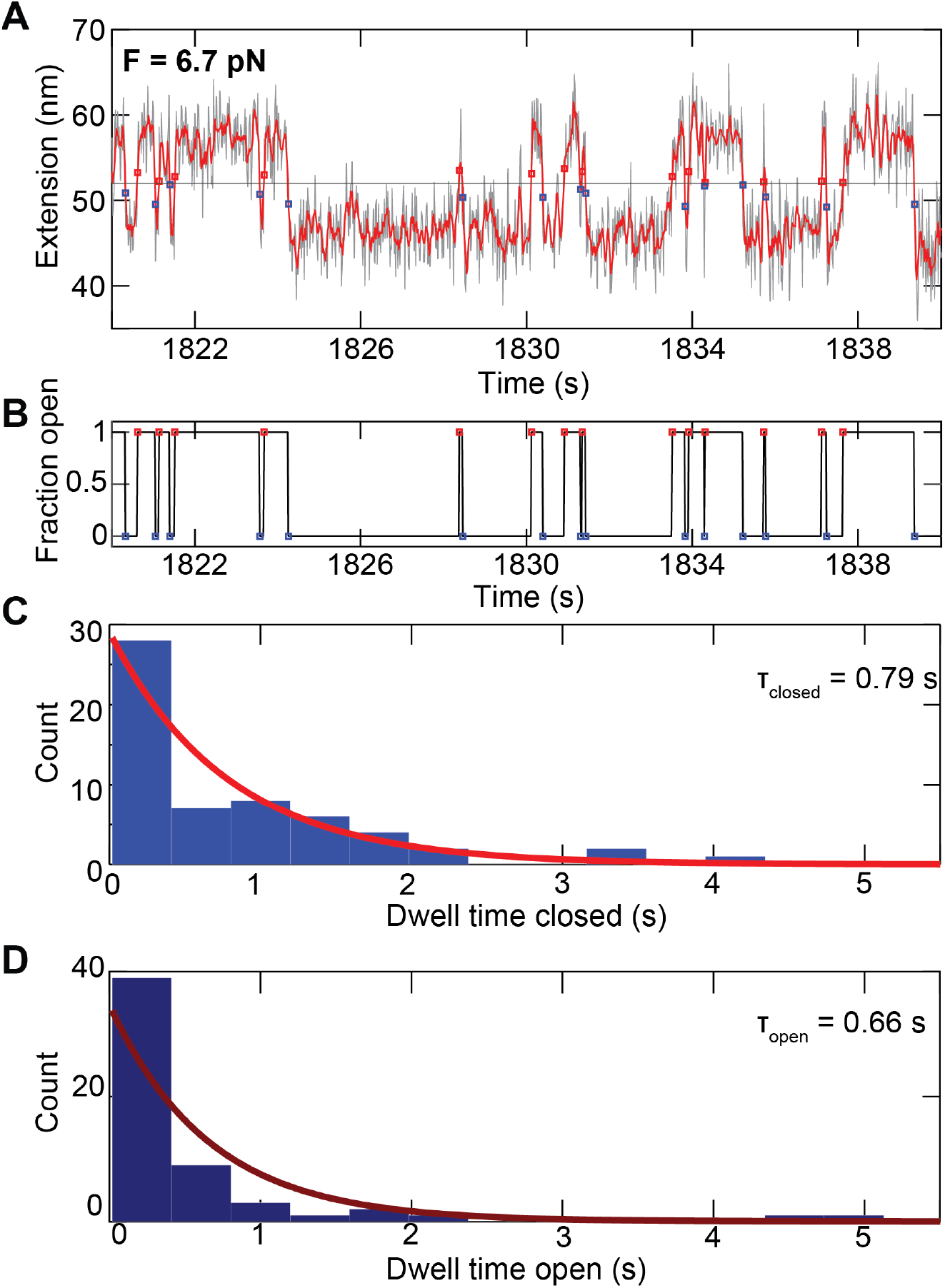
Example dwell time analysis of the tethered ligand extension time traces. **A** Short segment of an extension time trace measured for an ACE2-85 aa linker-SARS-CoV-2 RBD tethered ligand construct at a stretching force of 6.5 pN. Raw data at 58 Hz are shown in black and filtered data at 12 Hz in red. Assignment of the dwell times is based on the filtered data. The black horizontal line is the threshold; red squares indicate the first data point after crossing the threshold from below, i.e. transition from the closed to the open state; blue squares indicate the first data point after crossing the threshold from above, i.e. transition from the open to the closed state. **B** Time trace derived from the analysis shown in panel A, indicating the current state of the tether-ligand system with “1” corresponding to the open state and “0” to the closed state. The time between the transition between “0” and “1” correspond to the dwell times. **C, D** Histograms of dwell times in the closed state (**C**) and open state (**D**) obtained from the analysis shown in panels A and B (however using a longer trace, of which the data shown in A and B are just a subset). The dwell times are well described by single exponential fits, shown as solid line.

**Supplementary Table 1.** Equilibrium binding data for ACE2 binding to SARS-CoV-1 or SARS-CoV-2 RBD or S proteins. Studies for both ACE2 binding to RBD constructs and to the S protein are included; Wrapp et al. ^4^ find *K*_d_ = 14.7 nM for ACE2 binding to SARS-CoV-2 S and *K*_d_ = 34.6 nM for ACE2 binding to SARS-CoV-2 RBD, indicating similar affinities. Similarly, Yang et al. observe similar binding constants and mechanical stabilities for ACE2 binding to either the RBD or S using AFM force spectroscopy^5^.

| Study | ACE2 binding to SARS-CoV-1 RBD | ACE2 binding to SARS-CoV-2 RBD | Method and Comments |
| --- | --- | --- | --- |
| Lan et al.<sup>2</sup> | $K_d = 31$ nM<br>$k_{\text{sol,off}} = 4.3 \cdot 10^{-2} \text{ s}^{-1}$<br>$k_{\text{sol,on}} = 1.4 \cdot 10^6 \text{ s}^{-1} \text{ M}^{-1}$ | $K_d = 4.7$ nM<br>$k_{\text{sol,off}} = 6.5 \cdot 10^{-3} \text{ s}^{-1}$<br>$k_{\text{sol,on}} = 1.4 \cdot 10^6 \text{ s}^{-1} \text{ M}^{-1}$ | Surface-plasmon resonance |
| Shang et al.<sup>6</sup> | 185 nM<br>$k_{\text{sol,off}} = 3.7 \cdot 10^{-2} \text{ s}^{-1}$<br>$k_{\text{sol,on}} = 2.0 \cdot 10^5 \text{ s}^{-1} \text{ M}^{-1}$ | 44.2 nM<br>$k_{\text{sol,off}} = 7.8 \cdot 10^{-3} \text{ s}^{-1}$<br>$k_{\text{sol,on}} = 1.75 \cdot 10^5 \text{ s}^{-1} \text{ M}^{-1}$ | Surface-plasmon resonance |
| Starr et al.<sup>7</sup> | 0.12 nM | 0.039 nM | Yeast display screen |
| Walls et al.<sup>8</sup> | $K_d = 5.0 \pm 0.1$ nM<br>$k_{\text{sol,off}} = 8.7 \pm 5.1 \cdot 10^{-4} \text{ s}^{-1}$<br>$k_{\text{sol,on}} = 1.7 \pm 0.7 \cdot 10^5 \text{ s}^{-1} \text{ M}^{-1}$ | $K_d = 1.2 \pm 0.1$ nM<br>$k_{\text{sol,off}} = 1.7 \pm 0.8 \cdot 10^{-4} \text{ s}^{-1}$<br>$k_{\text{sol,on}} = 2.3 \pm 1.4 \cdot 10^5 \text{ s}^{-1} \text{ M}^{-1}$ | Bio-layer interferometry; uses S protein for both variants |
| Wang et al.<sup>9</sup> | $408 \pm 11$ nM<br>$k_{\text{sol,off}} = 1.9 \pm 0.4 \cdot 10^{-3} \text{ s}^{-1}$<br>$k_{\text{sol,on}} = 2.9 \pm 0.2 \cdot 10^5 \text{ s}^{-1} \text{ M}^{-1}$ | $95 \pm 7$ nM<br>$k_{\text{sol,off}} = 3.8 \pm 0.2 \cdot 10^{-3} \text{ s}^{-1}$<br>$k_{\text{sol,on}} = 4.0 \pm 0.2 \cdot 10^4 \text{ s}^{-1} \text{ M}^{-1}$ | Surface-plasmon resonance; uses S1 domain for SARS-CoV-2 |

